# CexE Is A Virulence Factor of *Citrobacter rodentium* And Present In Enteric Pathogens Of Humans

**DOI:** 10.1101/375527

**Authors:** Zachary Rivas, Ryan M. McCormack, Becky Adkins, George P. Munson

## Abstract

CexE is a 12 kDa protein that was originally reported to be present in just three strains of enterotoxigenic *Escherichia coli* (ETEC); a frequent causes of diarrheal illnesses worldwide. However, an examination of recently sequenced genomes has revealed that CexE is actually present in a majority of ETEC strains. Homologs of CexE are also present in enteroaggregative *E. coli* (EAEC) and other enteric pathogens including *Yersinia enterocolitica, Providencia alcalifaciens*, and *Citrobacter rodentium*. CexE and its homologs are expressed within virulence regulons of ETEC, EAEC, and *C. rodentium*. This, along with its distribution across several species of enteric pathogens, suggest that CexE confers a selective advantage to these pathogens. However, this hypothesis has yet to be tested *in vivo*. Here we demonstrate that CexE is conditionally secreted to the external leaf of the outer membrane of ETEC. Although CexE does not appear to play a role in adherence *in vitro*, it does facilitate colonization of murine intestinal tissues by *C. rodentium in vivo*. In adult mice wild-type bacteria reached significantly higher loads and were shed in higher numbers than a *cexE::kan* mutant. A similar trend was observed in neonatal mice. In addition, all of the neonates infected with the wild-type strain succumbed to infection within 16 days of inoculation. In contrast, 45% of the neonates infected with the *cexE::kan* strain survived for the 30 day duration of the experiment. These finding indicate that CexE is a conditionally secreted virulence factor that increases the colonization of hosts by enteric pathogens.

## Introduction

Approximately half a million infants and children perish from diarrheal diseases every year (1). For citizens of low–income countries diarrheal diseases are projected to remain among the ten leading causes of death through 2030 (2). Of the many viral and bacterial pathogens that cause diarrhea enterotoxigenic *Escherichia coli* (ETEC) is one of the most frequent causes with at least 280 million people sickened annually; most in low-income nations (3). ETEC causes diarrhea by elaborating enterotoxins that disrupt the normal flow of ions across the apical membranes of enterocytes. This disruption leads to a net loss of water into the lumen of the intestinal tract resulting in profuse watery diarrhea (4). In some instances ETEC induced diarrhea results in severe dehydration that can result in death if not adequately addressed.

Although enterotoxins are the penultimate cause of diarrhea, ETEC must also colonize the gastrointestinal tract to cause disease. Colonization is itself dependent upon the expression of pili that function as adherence factors facilitating the direct attachment of ETEC to enterocytes (5, 6). Although ETEC are heterogeneous and there are numerous pilus serotypes, many are positively regulated at the level of transcription by the ETEC virulence regulator Rns (CfaD) (7, 8). In vitro DNase I footprinting studies have shown that each Rns activated pilin promoter contains a Rns DNA binding site immediately upstream of the promoter’s −35 hexamer (9–11). In most cases the promoter proximal binding site is usually accompanied by one or more additional Rns binding sites further upstream. However site directed mutagenesis studies have shown that occupancy of the promoter proximal site has the greatest contribution to promoter activation (9, 11).

An in silico analysis of the genome of ETEC strain H10407 -a strain that has been used in several human challenge studies-led to the identification of two Rns binding sites upstream of *cexE* (12–14). As with pilin promoters, one of the binding sites was located immediately upstream of the −35 hexamer of *cexEp* and the other further upstream. As expected from the positions of its binding sites, Rns was found to activate the expression of *cexE*. Although *cexE* does not encode a pilin nor is it required for the expression of pili, its regulation by the virulence regulator Rns implies that *cexE* encodes a virulence factor. However options to test this hypothesis are limited by the fact that H10407 is strictly a human pathogen that does not naturally infect mice nor other laboratory animals. Fortunately a homolog of *cexE* is present in the genome of *Citrobacter rodentium* (15). Because this natural murine pathogen is a well-developed model for enteric infections, we used murine *C. rodentium* challenge studies to interrogate the function of CexE *in vivo* (16).

## RESULTS

### CexE is broadly distributed across ETEC strains

CexE was initially identified as a member of the Rns regulon by an in silico search of the genome of ETEC strain H10407 for Rns binding sites (14). Since no other ETEC genomes were publicly available at that time, it was unknown if CexE was restricted to a few isolates or more broadly distributed amongst ETEC strains. However in the intervening years over 200 ETEC genomes have been sequenced and deposited in NCBI’s databases; most as whole genome shotgun (WGS) sequences. Therefore to gauge the distribution of CexE across ETEC strains we ran iterative BLASTN searches of NCBI’s databases to identify genes with homology to *cexE*. After each search the most distal member was used as the query for a subsequent search until no additional members were found. BLASTP searches were not effective because automated annotation pipelines apparently have a high failure rate with regards to the identification of *cexE* alleles. The reasons for this are unclear as these genes are open reading frames of ca. 360 bp; most with recognizable Shine-Dalgarno sequences upstream of ATG start codons.

Of the 250 strains that were identified as ETEC –either explicitly in the record definition lines or by the presence of genes encoding the heat-labile (LT) and/or heat-stable (ST) enterotoxins– 76% were found to harbor *cexE* alleles. The prevalence of *cexE* alleles is similar to the genes encoding LT and ST which are present in 64% and 78% of strains identified as ETEC respectively. As with the enterotoxin genes, the prevalence of *cexE* alleles suggest a selective advantage for their distribution and maintenance amongst heterogeneous ETEC strains. The 190 alleles of *cexE* encode eleven distinct polypeptides (Fig. 1). Hereafter these will be distinguished from each other by Greek letter subscripts starting with CexEα that was first identified by Pilonieta et al. in 2007 (14). Sequences of each are presented in S. Fig. 1. The amino terminus of each polypeptide contains a putative signal peptide which suggest each polypeptide is transported to the periplasm (17, 18). Subsequent cleavage of the signal peptide from CexEα by a periplasmic signal peptidase has been confirmed experimentally (14).

**FIG 1.**
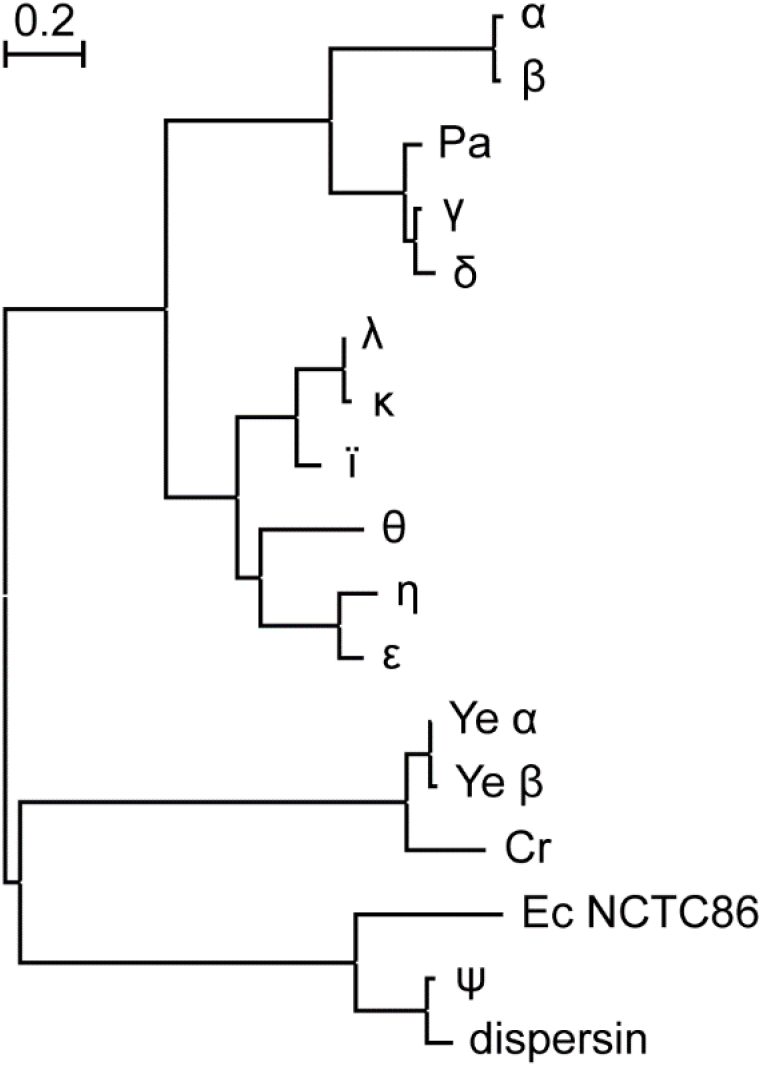
CexE homologs are prevalent amongst ETEC and a subset of other enteric pathogens. Proteins from ETEC strains are designated with a single Greek letter. Dispersin is from EAEC strain 042. The abbreviations Cr, Pa, and Ye indicate sequences found in *C. rodentium, P. alcalifaciens*, and *Y. enterocolitica* respectively. Ec NCTC86 is the strain of *E. coli* that was originally isolated by Theodor Escherich in 1885. The phylogram was constructed using the maximum likelihood method on a multiple sequence alignment generated by the MUSCLE algorithm; both of which were run within the MEGA7 software package (19).

With the lone exception of CexE_ψ_ all variants of CexE from ETEC strains clustered within the same clade. Within the CexE_ψ_ clade are two homologs that were found in 60% of the 162 *Yersinia enterocolitica* strains in NCBI’s WGS databases. In contrast to the situation in ETEC there is little variation between *Y. enterocolitica* strains; 93% of the polypeptides are identical to Ye_α_ found in a strain of *Y. enterocolitica* (ERL04757) that was isolated from human blood (20). Other species of *Yersinia* apparently lack CexE homologs. A CexE homolog is also present in the two complete genomes of *Citrobacter rodentium* as well as in one isolate of *Providencia alcalifaciens*. The former is a murine enteric pathogen while the latter has been described as an emerging enteric pathogen of humans (21–23). Notably the *P. alcalifaciens* homolog resides within a clade of ETEC proteins while the *Y. enterocolitica* and *C. rodentium* homologs are more closely related to dispersin of enteroaggregative *E.coli* (EAEC) (Fig. 1). Also within the dispersin clade is CexE_ψ_ from ETEC as well as a polypeptide from *E. coli* strain NCTC86. The latter was isolated by Theodor Escherich in 1885 as he sought to identify the cause of neonatal dysentery (24, 25).

### CexE_α_ is a conditional coat protein in ETEC

Dispersin of EAEC has an amino terminal signal peptide and enters the periplasm through the general secretory pathway (18, 26). In a second translocation event dispersin crosses the outer membrane but remains associated with the outer membrane as a coat protein through electrostatic interactions (26). To determine if CexE_α_ and CexE_Cr_ are also coat proteins proteinase K was used as a membrane impermeable probe of their subcellular distribution. Robust expression of the target proteins dictated the culture media that were used. When ETEC was cultured in CFA CexE_α_ was well expressed but was not digested by proteinase K unless the outer membrane was permeabilized by Triton/EDTA (Fig. 2A). Quantification of multiple immunoblots relative to DnaK –a cytoplasmic protein that was not degraded even when the outer membrane was permeabilized– revealed that there was no significant degradation of CexE_α_ in the absence of Triton/EDTA (S. Fig. 2A). Thus, CexE_α_ is localized in the periplasm when ETEC is cultured in CFA medium. Strikingly, proteinase K digested > 98% of CexE_α_ when ETEC was grown in IMDM even in the absence of outer membrane permeabilization (Fig. 2A, S. Fig. 2A). A western blot of TCA precipitated IMDM supernatants found no detectable CexE_α_ in the extracellular milieu (S. Fig. 3). These results demonstrate that translocation of CexE_α_ across the outer membrane is influenced by culture conditions and that when it is secreted it remains predominately associated with the outer membrane. In contrast, CexE_Cr_-FLAG was not translocated across the outer membrane when *C. rodentium* was cultured in IMDM nor LB (Fig. 2B). It is unlikely that the < 3 kDa FLAG epitope tag prevented the translocation of CexE_Cr_ because fusion of the 30 kDa TEM-1 β-lactamase enzyme to the carboxy terminus of CexE_α_ did not prevent its translocation across the outer membrane of ETEC (S. Fig. 4). Given the conditional secretion of CexE_α_ a more plausible explanation for the absence of CexE_Cr_ translocation is that the in vitro conditions that were tested were not conducive to its translocation.

**FIG 2.**
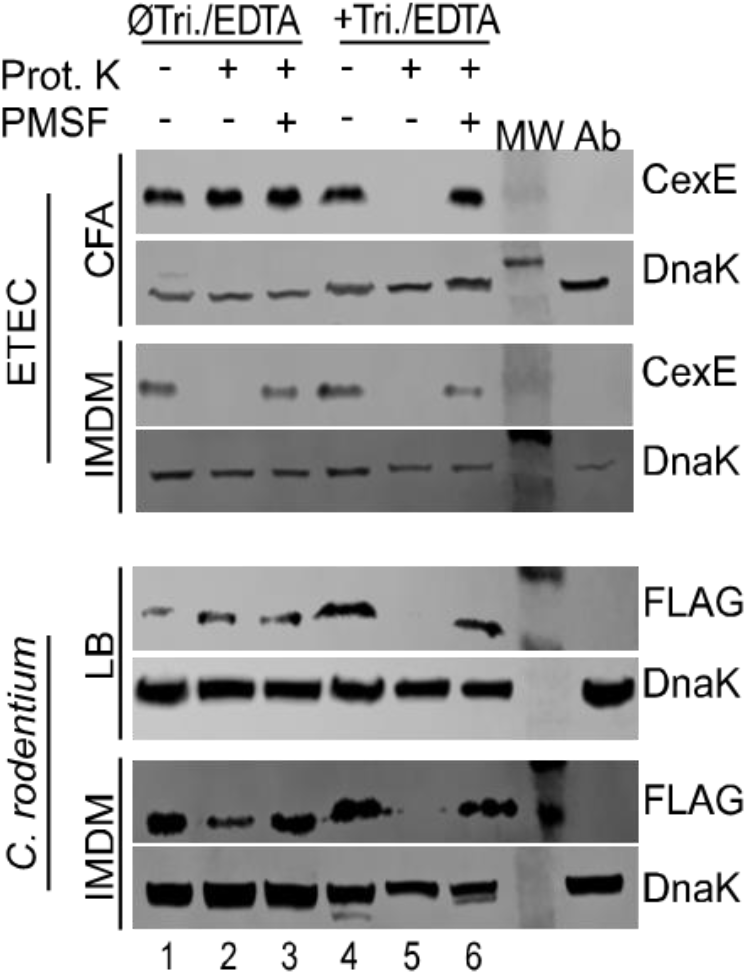
Culture medium influences the subcellular distribution of CexE_α_. Western blots of whole cell lysates of ETEC (H10407) or *C. rodentium* (GPM1830) in which the outer membrane was either left intact or permeabilized with Triton X-100 and EDTA prior to addition of proteinase K. Permeabilization was required for the digestion of CexE_α_ when ETEC was grown in CFA but not IMDM. This indicates that CexE_α_ is exposed on the surface of ETEC when the pathogen is cultured in the latter medium but remains within the bacterial envelope when ETEC is cultured in CFA. Likewise FLAG tagged CexE_Cr_ was retained within the envelope of *C. rodentium* when the murine pathogen was cultured in LB. Unlike with ETEC, IMDM did not result in translocation of CexE_Cr_ across the outer membrane. Lane numbering corresponds to quantification presented in S. Fig. 2. Abbreviations: Tri., Triton X-100; Prot. K, proteinase K; MW, molecular weight ladder; Ab, antibody specificity controls. Antibody specificity control lanes were loaded with whole cell lysates of ETEC strain GPM1163 (***cexE**_α_**::kan***) or ***C. rodentium*** strain DBS100 which do not produce CexE_α_ or CexE_Cr_-FLAG respectively.

### CexE_α_ and CexE_Cr_ do not significantly modulate adherence

After finding that CexE_α_ can associate with the outer face of the outer membrane we next sought to determine if it has an effect on adherence. Although we observed that a *cexE_α_::kan* mutant was slightly more adherent to HCT-8 cells than WT ETEC, the difference was not statistically significant (Fig. 3A). Likewise there was no significant different between WT *C. rodentium* and a *cexE*_Cr_ mutant (Fig. 3B). However, the latter result was expected because CexE_Cr_ is not a coat protein under these conditions (Fig. 2B). In contrast, an insertion in *rns* –which abolishes expression of both CexE_α_ and the adhesive CFA/I pilus– significantly decreased adherence of the mutant compared to both the WT and *cexE_α_::kan* strains of ETEC. These results suggest that CexE_α_ does not significantly modulate the adherence of ETEC even when the pathogen is cultured in IMDM; a condition in which CexE_α_ coats the outer membrane.

**FIG 3.**
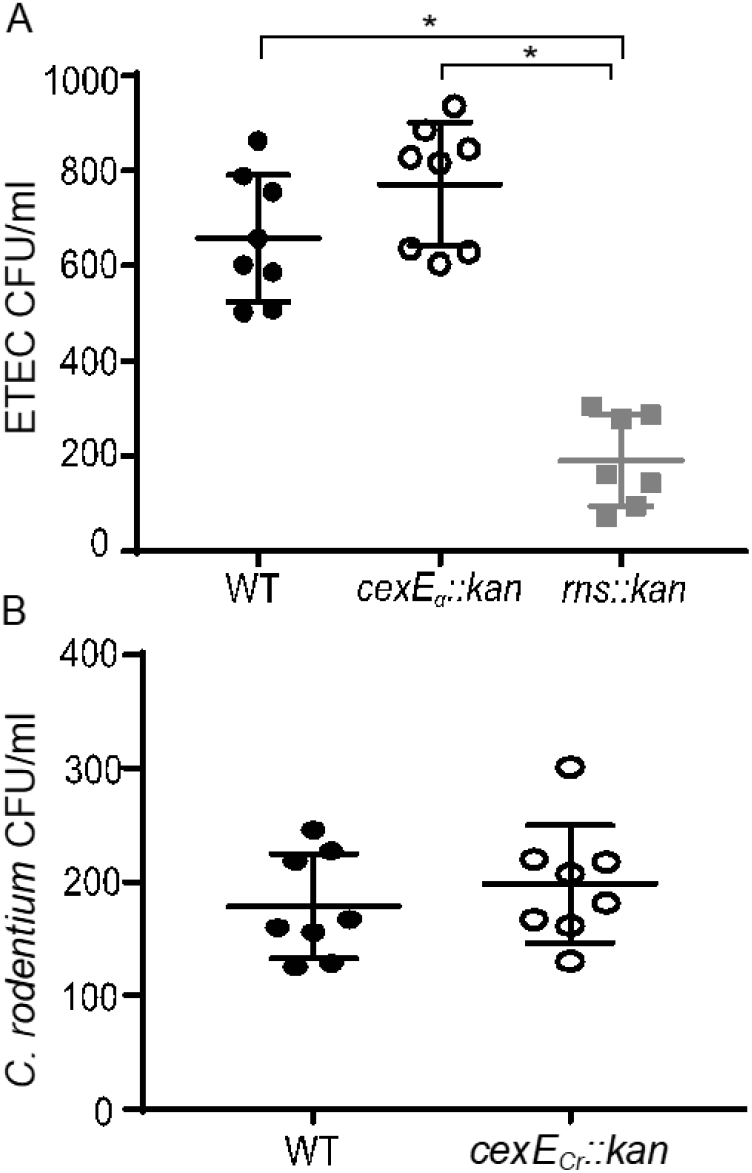
CexE_α_ and CexE_Cr_ do not significantly modulate adherence to mammalian cells. Adherent bacteria were enumerated after a 1 hr infection of HCT-8 cells in IMDM 10% FBS. **(A)** The difference between WT ETEC (GPM1168) and a *cexE_α_::kan* (GPM1163) mutant was not statistically significant. In contrast, a mutant (GPM1236) lacking the Rns virulence regulator was significantly impaired in its ability to adhere to HCT-8 cells. **(B)** As with ETEC strains, the difference between WT *C. rodentium* (GPM1831) and a *cexE*_Cr_*::kan* mutant (GPM1827) was not statistically significant. \**P* < 0.0001 by Tukey’s multiple comparisons test.

### CexE_Cr_ is a virulence factor *in vivo*

Although we did not observe secretion of CexE_Cr_ *in vitro* we considered it plausible that CexE_Cr_ is a virulence factor because its expression is regulated by the virulence regulator RegA (27). Its presence in a natural murine pathogen is also fortuitous because there is no small animal model to evaluate the function of dispersin in EAEC. In addition the murine ETEC model has several limitations due to the fact that ETEC does not normally colonize mice (28). Thus we chose *C. rodentium* for our *in vivo* studies because it is the most appropriate of the three pathogens for murine infection studies. To determine whether or not CexE_Cr_ is a virulence factor we orogastrically inoculated C57BL/6 mice with *C. rodentium* strain GPM1831a or GPM1827 (*cexE_Cr_::kan*). GPM1831a carries a kanamycin cassette in an intergenic region downstream of *cexE* and was considered our WT strain. On each of the three selected days after inoculation the infected animals shed fewer *cexE_Cr_::kan* than WT bacteria (Fig. 4). Although the difference was not statistically significant on day three it was significant on days six and ten post inoculation. In addition mice shed greater numbers of both strains as the infection progressed. For WT *C. rodentium* this was expected based on previous *in vivo* studies (16). These results were not restricted to C57BL/6 mice as similar results were observed when 129X1/SvJ mice were used (S. Fig. 5). Not surprisingly, the differences in fecal shedding reflected the differences between the two strains within the intestines (Fig. 5).

**FIG 4.**
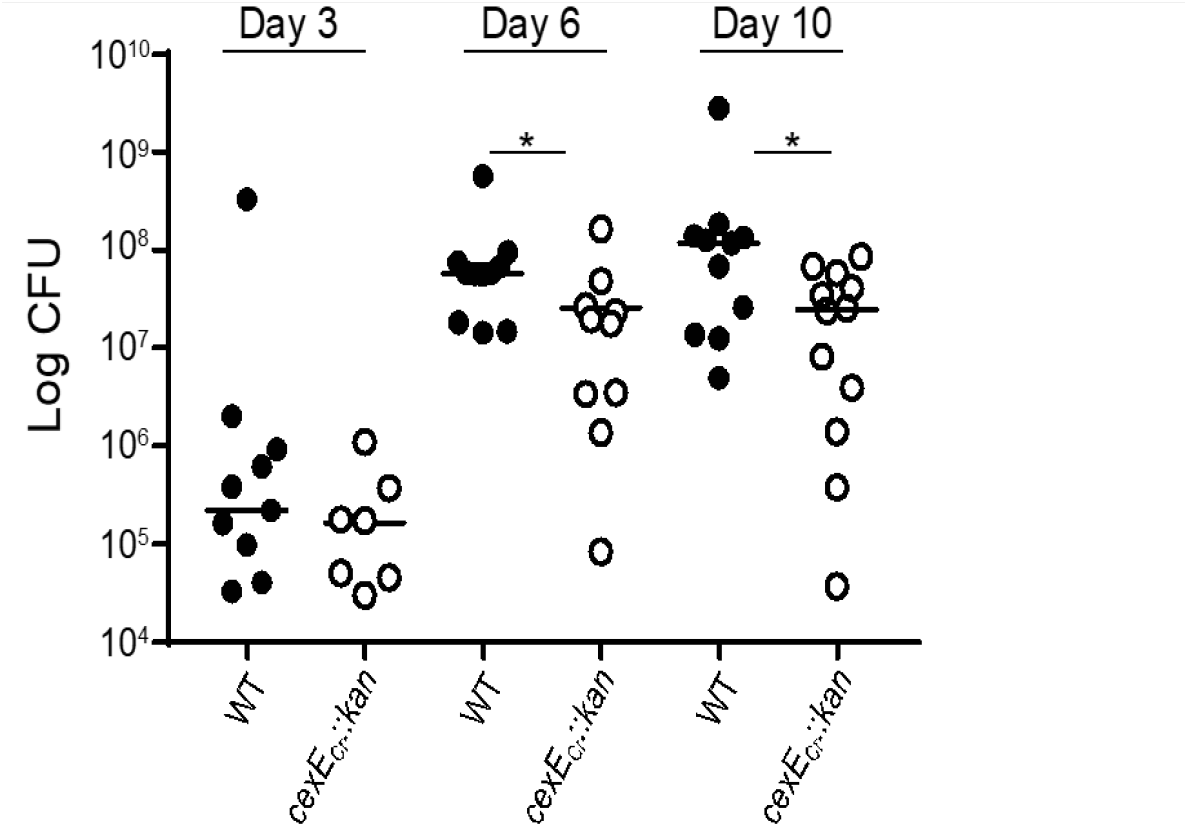
Fecal shedding of WT *C. rodentium* exceeds that of a *cexE_Cr_* mutant. C57BL/6 mice were orogastrically inoculated with 10^8^ CFUs with either WT (GPM1831a) or a *cexE_Cr_::kan* (GPM1827) mutant. Fecal pellets were collected on days 3, 6, and 10 post inoculation and CFUs were normalized to the mass of each pellet. Circles represent fecal pellets of separate animals; *n* ≥11 mice per group. \**P* < 0.05 by Mann-Whitney U test.

**FIG 5.**
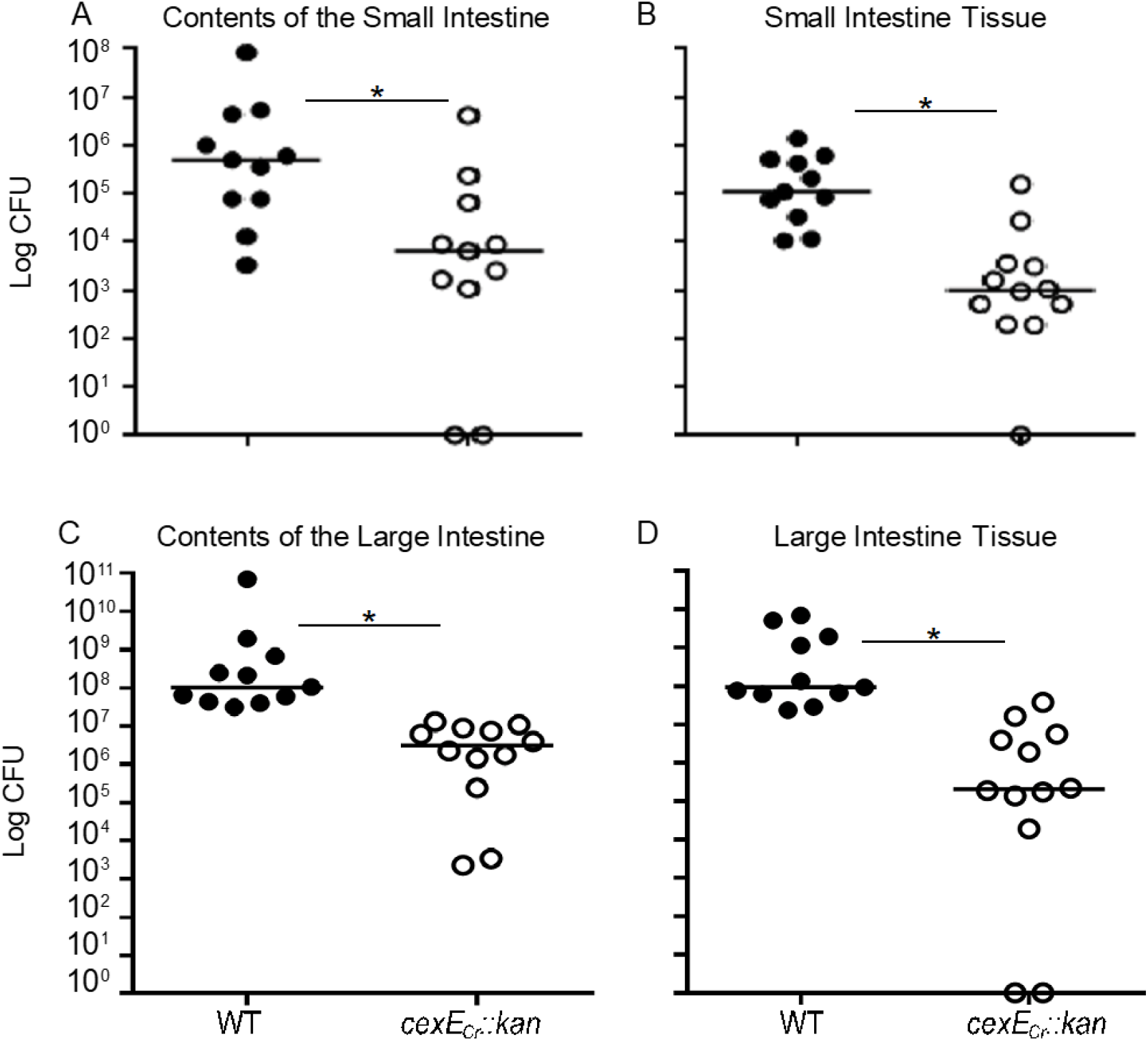
Intestinal loads of mice infected with a *cexE_Cr_* mutant are lower than those infected with WT *C. rodentium*. C57BL/6 mice were orogastrically inoculated with 10^8^ CFU of either WT *C. rodentium* (GPM1831a) or a *cexE_Cr_::kan* mutant (GPM1827). Tissues were harvested 10 days after inoculation. CFUs were normalized to sample mass. The differences between the two strains of bacteria were significant in both the **(A-B)** small intestine and its contents as well as in the **(C-D)** large intestine and its contents. \**P* < 0.005 by Mann-Whitney U test, n ≥11 mice per group.

To determine if the effect of CexE_Cr_ was age-dependent or -independent we also inoculated infant C57Bl/6 mice with both strains of *C. rodentium*. As with adult animals the loads of both strains increased as the infection progressed and significantly more WT than mutant bacteria were recovered from the large intestines of infant mice; for technical reasons the small intestines were not analyzed (Fig. 6). Although immunocompetent adult mice are able to clear *C. rodentium* infections, all infant mice infected with the WT strain perished (Fig. 6)(29). In contrast a significant number of the mice infected with the *cexE_Cr_::kan* strain survived for the duration of the experiment. In aggregate these results demonstrate that *C. rodentium* uses CexE_Cr_ to achieve higher intestinal burdens and increase its virulence in both adult and infant mice.

**FIG 6.**
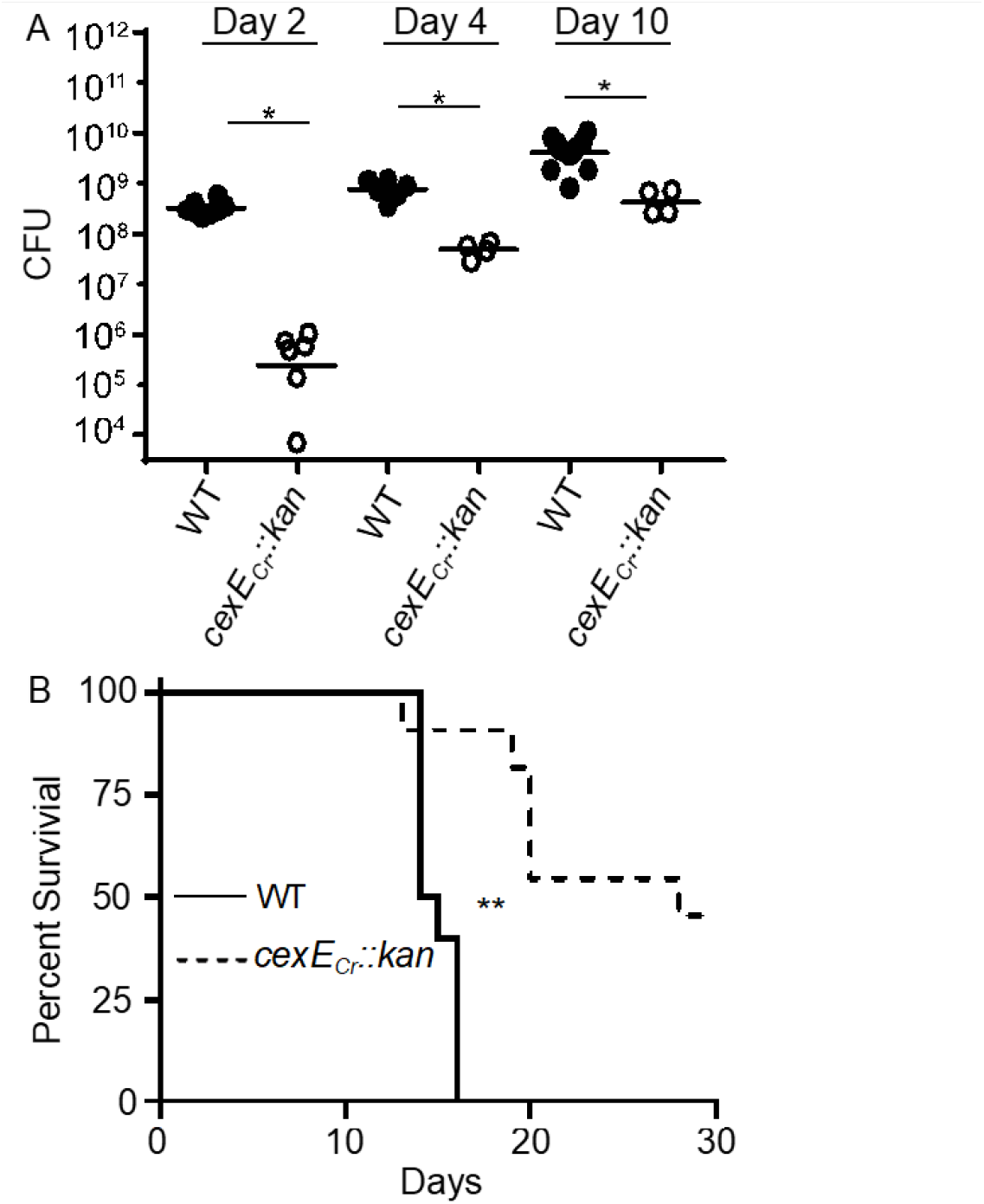
Infant mice infected with mutant *C. rodentium* had reduced colonization of their large intestines and enhanced survival compared to those infected with WT *C. rodentium*. Fifteen-day old infant C57Bl/6 mice were infected with 10^7^ CFU of WT *C. rodentium* (GPM1831a) or a *cexE_Cr_::kan* mutant (GPM1827). **(A)** *C. rodentium* colon titers, normalized to tissue mass, were determined at 2, 4, and 10 days post inoculation. * *P* < 0.05 \*\**P*<0.005 by Mann-Whitney U test. **(B)** In a parallel group of animals survival was monitored out to 30 days post inoculation. ≥ 10 animals were analyzed in two separate experiments. *** *P* = 0.0005 by log rank (Mantel-Cox) test.

## Discussion

A given isolate of *E. coli* that causes diarrheagenic disease in humans is often categorized as one of five pathovars; enterotoxigenic *E. coli* (ETEC), enteroaggregative *E. coli* (EAEC), enterohemorrhagic *E. coli* (EHEC), enteropathogenic *E. coli* (EPEC), and enteroinvasive *E. coli* (EIEC). A number of characteristics are used to distinguish these pathovars from one another including disease etiology, patterns of adherence, and repertoires of virulence genes. However this classification system belies the fact that isolates within each category may display considerable heterogeneity and originate from different phylogenetic lineages (30–32). Heterogeneity is particularly prevalent amongst ETEC isolates which is often considered an evolving pathogen. Thus, it is not surprising that although a majority of ETEC strains carry *cexE* alleles a minority do not. However, the fact that 76% of ETEC genomes harbor a *cexE* allele suggest a selective advantage to the gene’s acquisition and maintenance. Moreover, *cexE* homologs have been acquired by another *E. coli* pathovar –EAEC– as well as three other species of enteric pathogens; *P. alcalifaciens, Y. enterocolitica* and *C. rodentium*. Overall these observations suggest that CexE and its homologs contribute to the pathogenicity of some gram-negative enteric pathogens.

ETEC, EAEC, and *P. alcalifaciens* do not naturally infect mice; therefore, we considered them unsuitable to evaluate the role of CexE and its homologs in vivo. In contrast both *Y. enterocolitica* and *C. rodentium* are capable of infecting mice including when administered orogastrically; arguably the most relevant route for enteric pathogens. Of the latter two pathogens we chose *C. rodentium* for our in vivo studies for the simple reason that we were unable to obtain an isolate of *Y. enterocolitica* that harbors a *cexE* homolog. We found that mice inoculated with WT *C. rodentium* shed significantly more bacteria on days 6 and 10 post inoculation than mice inoculated with the *cexE_Cr_* mutant. The observed CexE_Cr_-dependent enhanced fecal shedding would seem to be consistent with the proposed function of the CexE homolog dispersin in EAEC. *In vitro* dispersin has been shown to decrease the adherence of EAEC to HEp-2 cells as well as human intestinal explants (26). Thus, as its name implies dispersin has been proposed to contribute to the pathogenecity of EAEC by increasing the dispersal of the pathogen throughout the intestinal tract. Decreased adherence would also be expected to result in greater fecal shedding of WT bacteria than *cexE* mutants. However, this model is at odds with the loads of *C. rodentium* within the intestinal tissues. In both the small and large intestines of adult mice, we found significantly higher loads of WT *C. rodentium* than the *cexE_Cr_* mutant. A similar pattern was also observed in infant mice. Although we did observe that WT ETEC are less adherent to HCT-8 cells than a *cexE* mutant, the difference was not significant. Therefore, greater shedding of the WT strain compared to the *cexE_Cr_* mutant is likely a consequence of higher loads of the WT strain within the intestines rather than an increased tendency of the WT strain to detach and disperse. Although the mechanism is currently unknown, our *in vivo* studies clearly demonstrate that CexE promotes colonization and/or survival of an enteric pathogen within the intestinal tract.

All CexE variants and dispersin have predicted amino terminal signal peptides that would result in their transport to the periplasm via the general secretory pathway (17). Such signal peptides are cleaved upon entry into the periplasm and cleavage of CexE_α_’s signal peptide has been experimentally verified (14). This is apparently followed by a second translocation event that transports CexE_α_ to the external face of the outer membrane that is dependent upon by environmental conditions such as growth medium. For dispersin of EAEC this second translocation event requires five genes, *aatPABCD* (33). AatA likely forms a conduit through the outer membrane because it is a homolog of TolC; a protein that has been shown to form β-barrel pores in the outer membrane of *E. coli* and other gram-negative bacteria (34, 35). The other *aat* genes encode essential accessory proteins because deletion of any of the five genes abolishes translocation of dispersin across the outer membrane (33). Homologs of all five genes are also present in ETEC strains harboring *cexE* as well as *C. rodentium, Y. enterocolitica*, and *P. alcalifaciens* (unpublished observation). This suggest that the mechanism of CexE/dispersin translocation across the outer membrane is conserved across pathovars and species. Although we did not observe translocation of CexE_Cr_ in vitro, this is probably due to an inability to replicate the appropriate conditions. Conditional translocation of CexE_Cr_ is a valid possibility because we observed that CexE_α_ was translocated across the outer membrane when ETEC was cultured in IMDM but not CFA medium. The mechanism of this conditional translocation is currently under investigation.

Dispersin has been shown to decrease the adherence of EAEC to enterocytes in vitro by coating the outer membrane of the pathogen (26). Although we observed that CexE_α_ also coats the outer membrane of ETEC, WT ETEC was no less adherent to HCT-8 cells than a *cexEα* mutant. Thus, CexE of ETEC does not appear to act in a dispersin-like manner which raises the question as to whether or not it acts locally at the outer membrane. The reported association of CexE_α_ with outer membrane vesicles (OMVs) presents an alternative hypothesis in which CexE_α_ acts distally upon host cells OMVs are bacteria-derived nanostructures that can deliver virulence factors to host cells and other targets (36, 37). For example, OMVs have been shown to contain and also be coated with the LT enterotoxin of ETEC (38). Such LT laden OMVs facilitate the delivery of LT to the cytosol of mammalian cells (38). Since CexE_α_ colocalizes with LT in OMVs and OMVs deliver LT to host cells it is plausible that CexE also reaches host cells. Therefore, unraveling the mechanism of CexE’s contribution to pathogenicity will require consideration that it may function locally at the bacterial outer membrane or alternatively at distal sites upon or within host cells.

## ACKNOWLEDGEMENTS

Research reported in this publication was supported by NIAID of the National Institutes of Health under award number R21AI128164. The contents of this publication are solely the responsibility of the authors and do not necessarily represent the official views of the NIH.

## SUPPLEMENTARY MATERIAL

**FIG S1.**
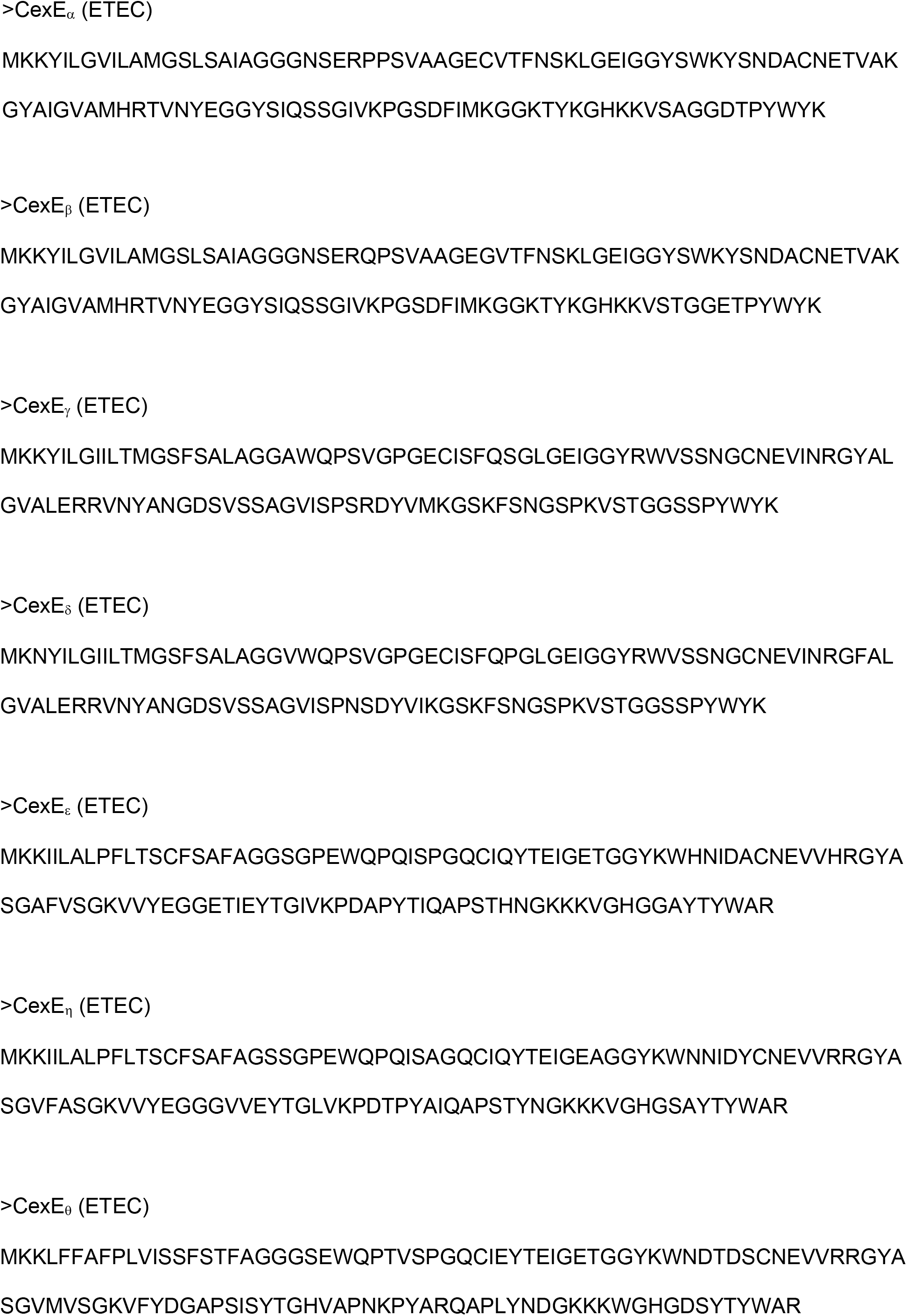
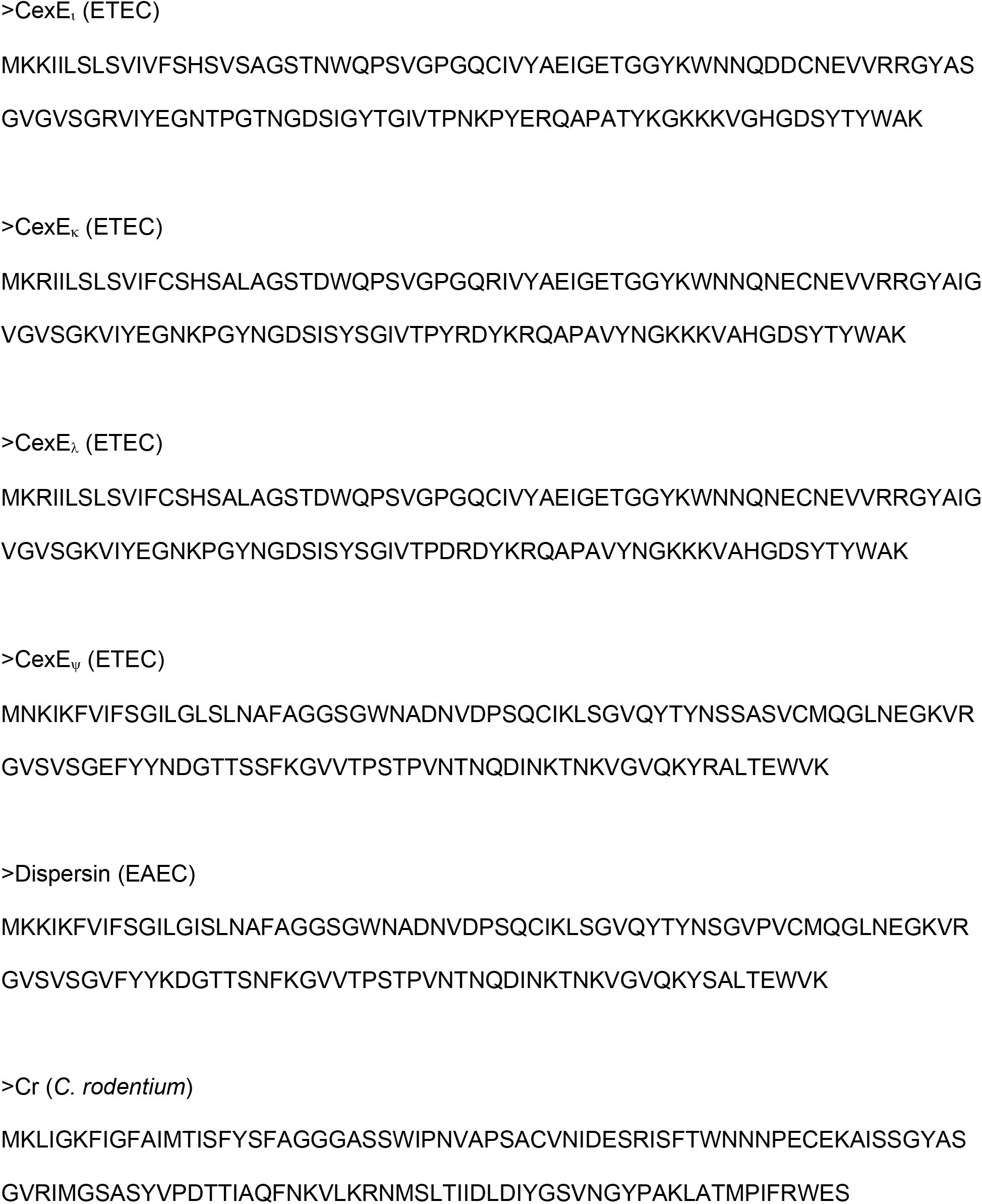
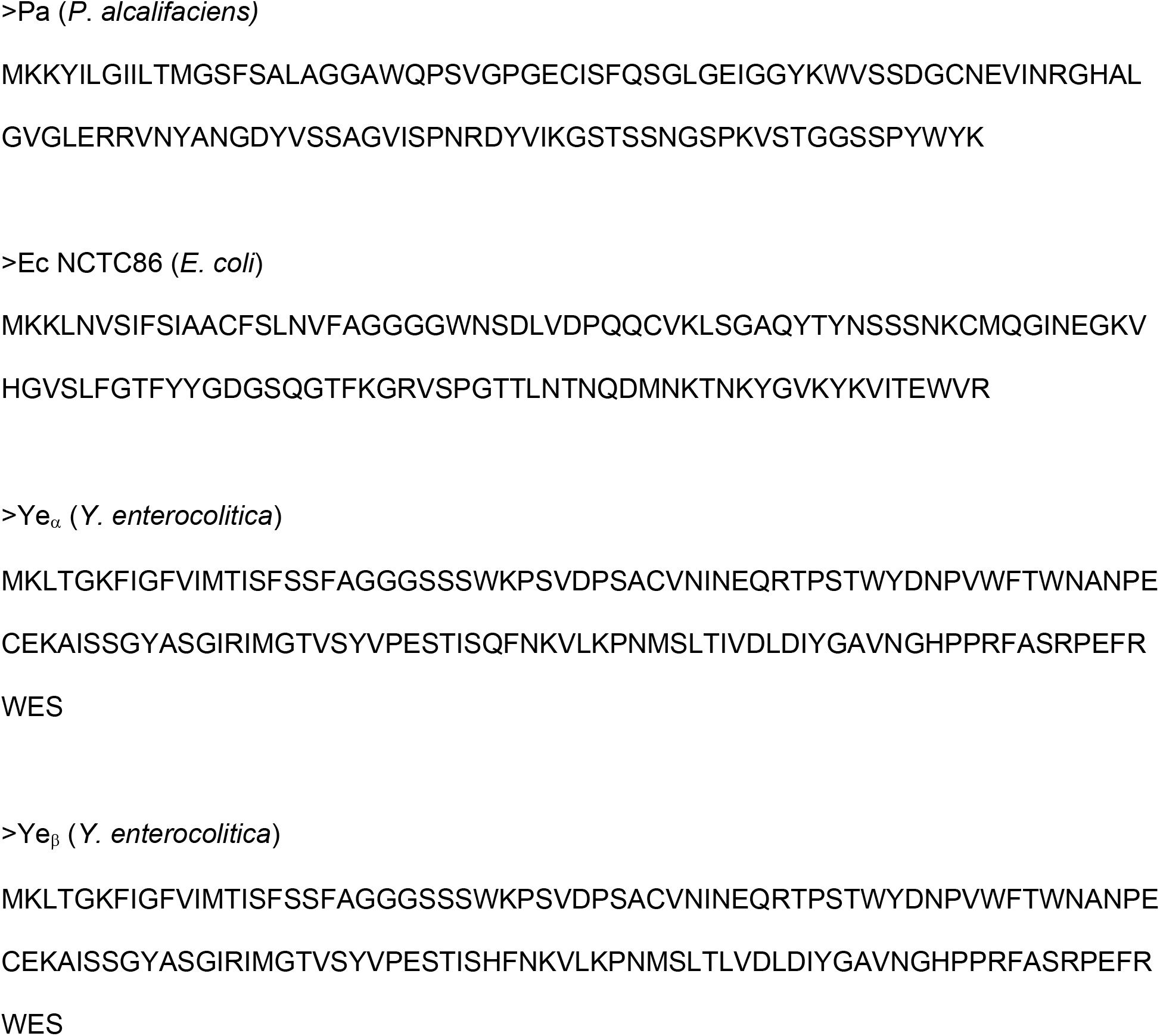
Sequences of CexE homologs

**FIG S2.**
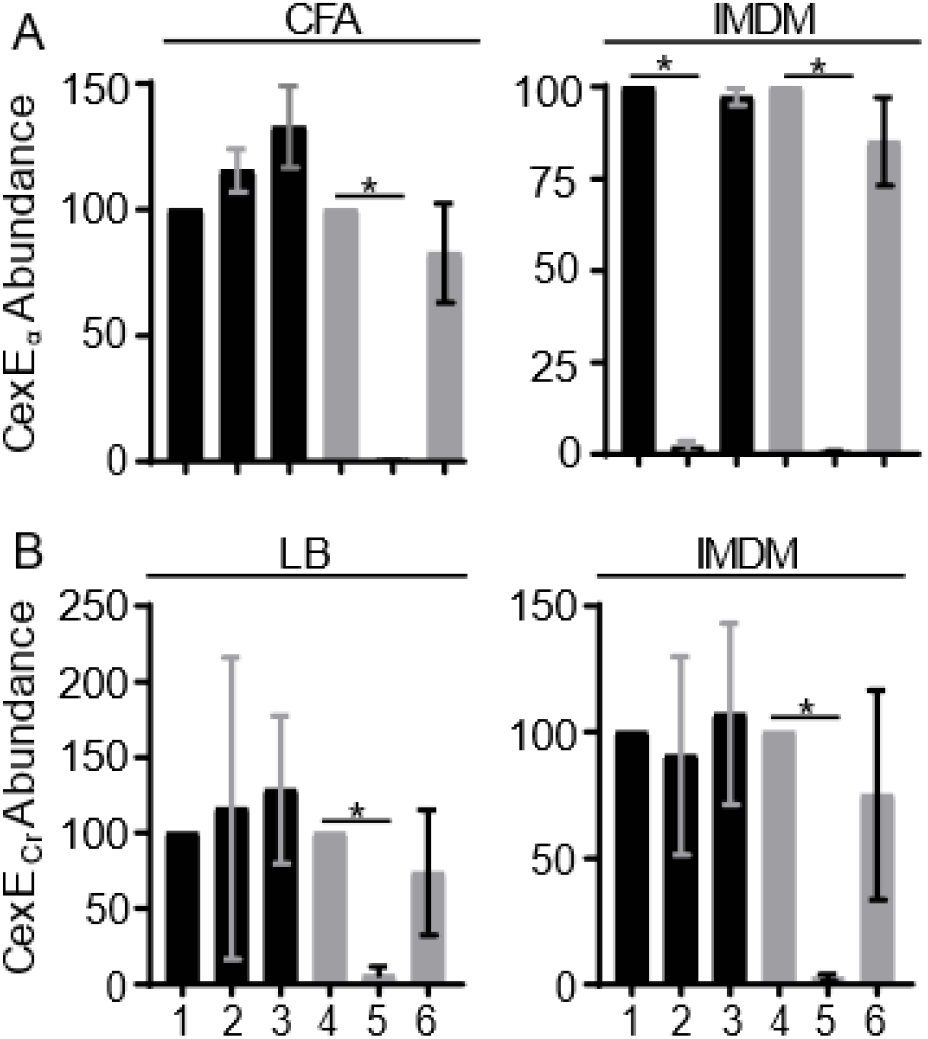
Quantification of proteinase K digestion of antigens relative to DnaK. **(A)** Quantification of CexE_α_ digestion from ETEC cultured in CFA or IMDM. **(B)** Quantification of CexE_Cr_ digestion from *C. rodentium* cultured in LB or IMDM. Column numbering corresponds to Western blot lanes in Fig 2. *n* = 3, \**P* < 0.0001 by Student’s *t*-test. Black columns correspond to untreated cells (PBS). Gray columns correspond to cells treated with PBS, Triton (1%), and EDTA (10mM).

**FIG S3.**
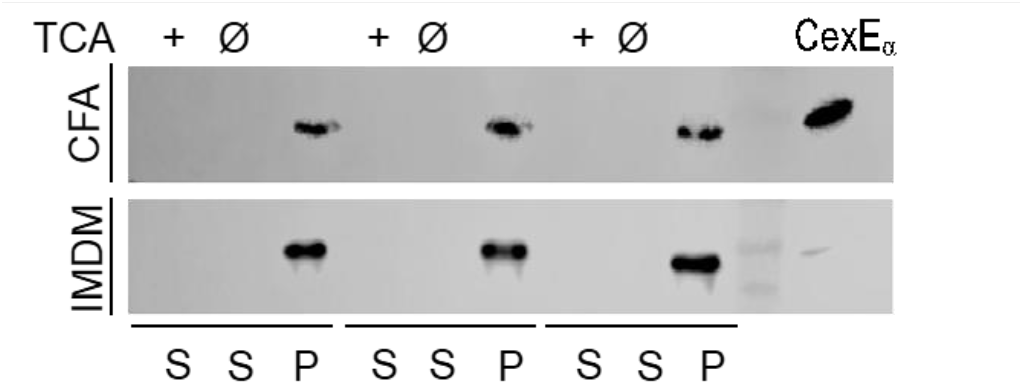
CexE_α_ is not detectable in the culture supernatants of ETEC. TCA precipitation of the supernatant of H10407 grown in either CFA or IMDM. Purified CexE_α_ was TCA precipitated as a positive control at approximately 10 ng of TCA precipitated protein. The limit of detection is estimated to be 10 ng of CexE_α_ Abbreviations: S, culture supernatant; P, cell pellet.

**FIG S4.**
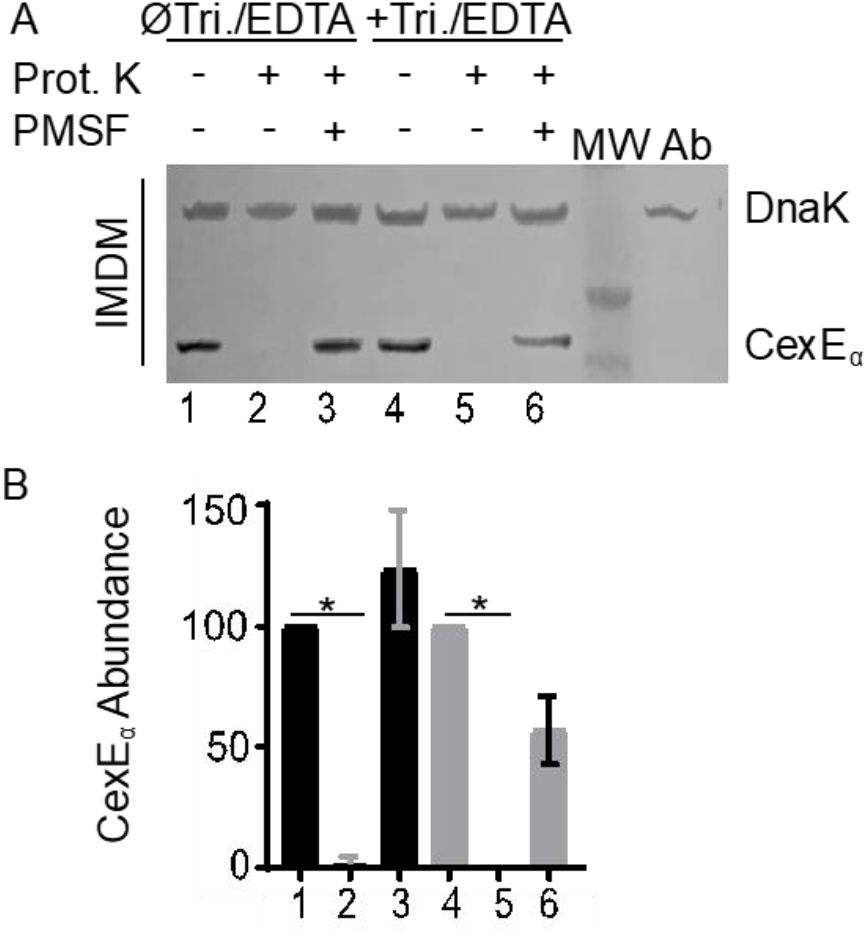
Translocation of CexE_α_-TEM-1 to the surface of ETEC cultured in IMDM. (A) Western blot of whole cell lysates of ETEC strain GPM1163 / pGPM1034bla probed with anti-CexE and DnaK antibodies. Membrane permeabilization is not required for enzymatic digestion of a 43 kDa CexE_α_-TEM-1 fusion protein indicating that the fusion protein is exposed on the surface of ETEC. (B) Quantification of CexE_α_-TEM-1 digestion relative to DnaK. Column numbers corresponds to (A) Western blot lane numbers. *n* = 3, \**P* < 0.0001 by Student’s t-test. Abbreviations: Prot. K, proteinase K; MW, molecular weight ladder; Ab, antibody specificity control with whole cell lysates of GPM1163 lacking the CexE_α_-TEM-1 fusion protein.

**FIG S5.**
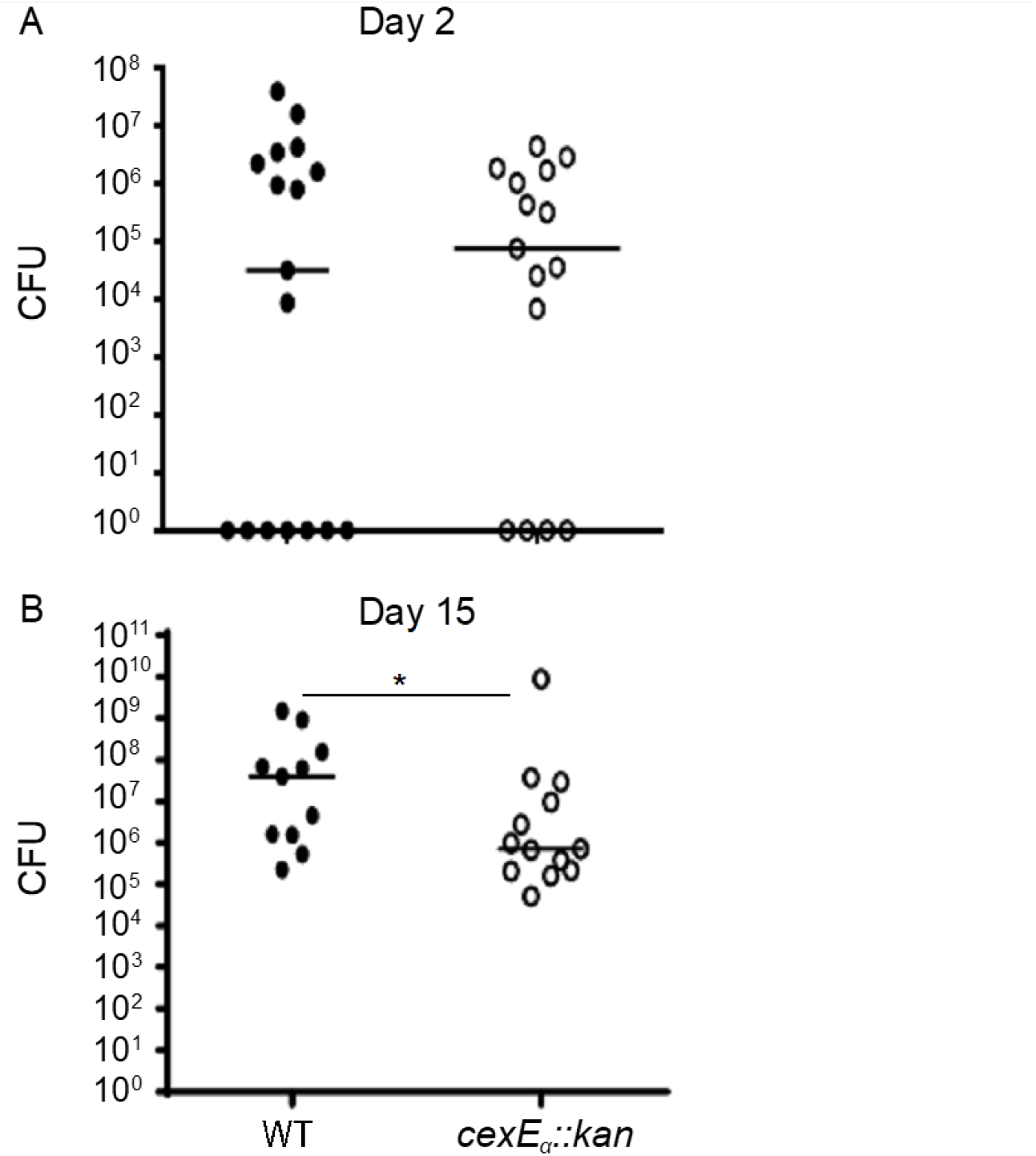
A CexE_Cr_ mutant has the same phenotype in 129X1/SJV and C57BL/6 mice. 129X1/SJV mice were orogastrically inoculated with 10^10^ CFUs with either WT *C. rodentium* (GPM1831a) or a *cexE_Cr_::kan* (GPM1827) mutant. Fecal pellets were collected on days 2 and 15 post inoculation and CFUs were normalized to the mass of each pellet. Although the difference between fecal shedding of the two strains was not significant at the early (A) time point it was statistically significant (B) 15 days after inoculation. Circles represent fecal pellets of separate animals; *n* ≥10 mice per group. \**P* <0.05 by Mann-Whitney U test

## MATERIALS AND METHODS

### Bacterial growth conditions

Bacteria were cultured aerobically at 37°C in CFA medium (39), Lysogeny Broth (LB; HIMEDIA), or Iscove’s Modified Dulbecco’s Medium (IMDM; Thermo Fisher Scientific). Growth media were supplemented with 50 μg/ml kanamycin or 100 μg/ml ampicillin as appropriate.

### Bacterial strains and plasmids

Plasmid pGPM1034bla expresses a CexE-TEM-1 fusion protein from *cexEp* or T7p. It was constructed by amplification of the β-lactamase gene from pUC19 with oligonucleotide primer pair 1126/1127. The PCR product was digested with XhoI then ligated into the same site of pGPM1034. λRED mediated recombineering was used for the construction of the following strains as previously described (40, 41). Kanamycin resistance cassettes targeting *cfaD* and *cexE_α_* were amplified from pKD4 with primer pairs 608/625 and 622/623 respectively. A cassette for the disruption of the *lac* operon was amplified from JF876 with primers 108/518. Electroporation of the cassettes into H10407 / pSIM6 resulted in recombinants GPM1236, GPM1163, and GPM1168 respectively. Cassettes for epitope tagging, disruption, or insertion downstream of *cexE_Cr_* were amplified from pSUB11 with primer pairs 1178/1179, 1177/1179, and 1193/1179 respectively. Electroporation of the cassettes into DBS100 / pSIM6 resulted in recombinants GPM1830, GPM1827, and GPM1831a respectively. Recombinants were cured of λRED expression plasmids by passage at 42°C. Insertions were verified by PCR with primers flanking the insertion sites.

**Table 1.**
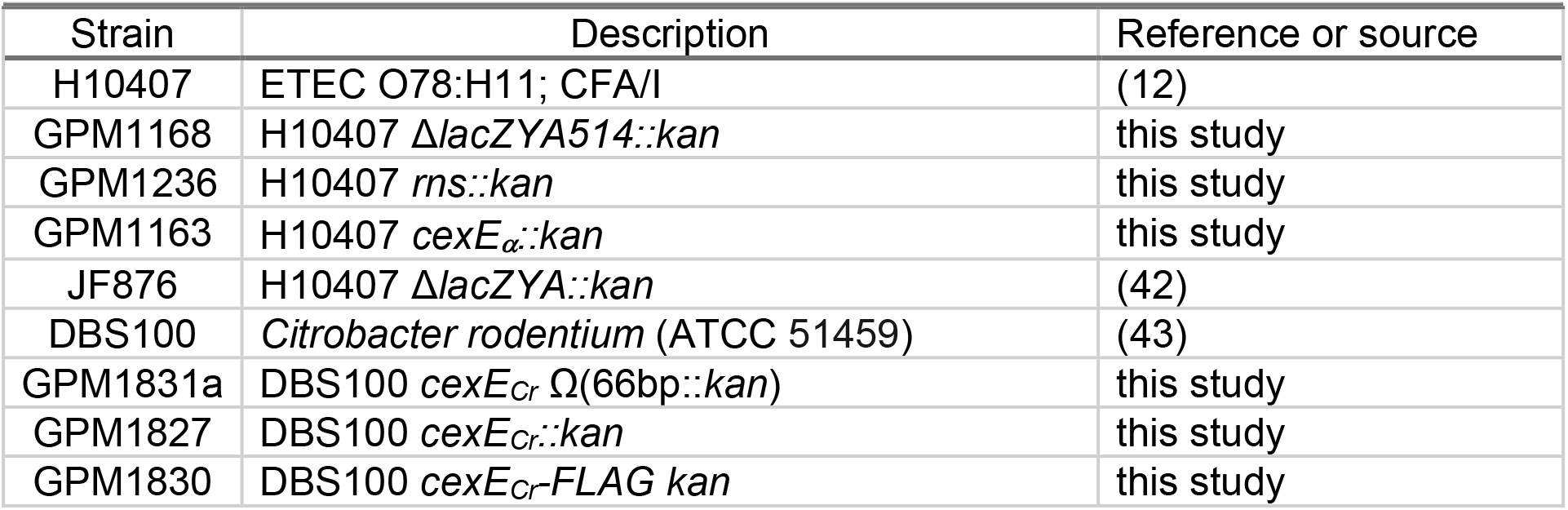
Strains used in this study

**Table 2.** Plasmids used in this study

| Name | Description | Marker | Reference or source |
| --- | --- | --- | --- |
| pNEB193 | cloning vector | Amp | New England Biolabs |
| pUC19 | cloning vector | Amp | (44) |
| pGPM1034bla | CexE-TEM-1 expressed from T7p or *cexEp* | Amp, Kan | this study |
| pGPM1034 | CexE expressed from T7p or *cexEp* | Kan | (14) |
| pKD4 | PCR template for *kan* cassettes | Amp, Kan | (40) |
| pSIM6 | λRED expression plasmid | Amp | (41) |
| pSUB11 | PCR template for 3XFLAG epitope tagging by λRED | Kan | (45) |

**Table 3.**
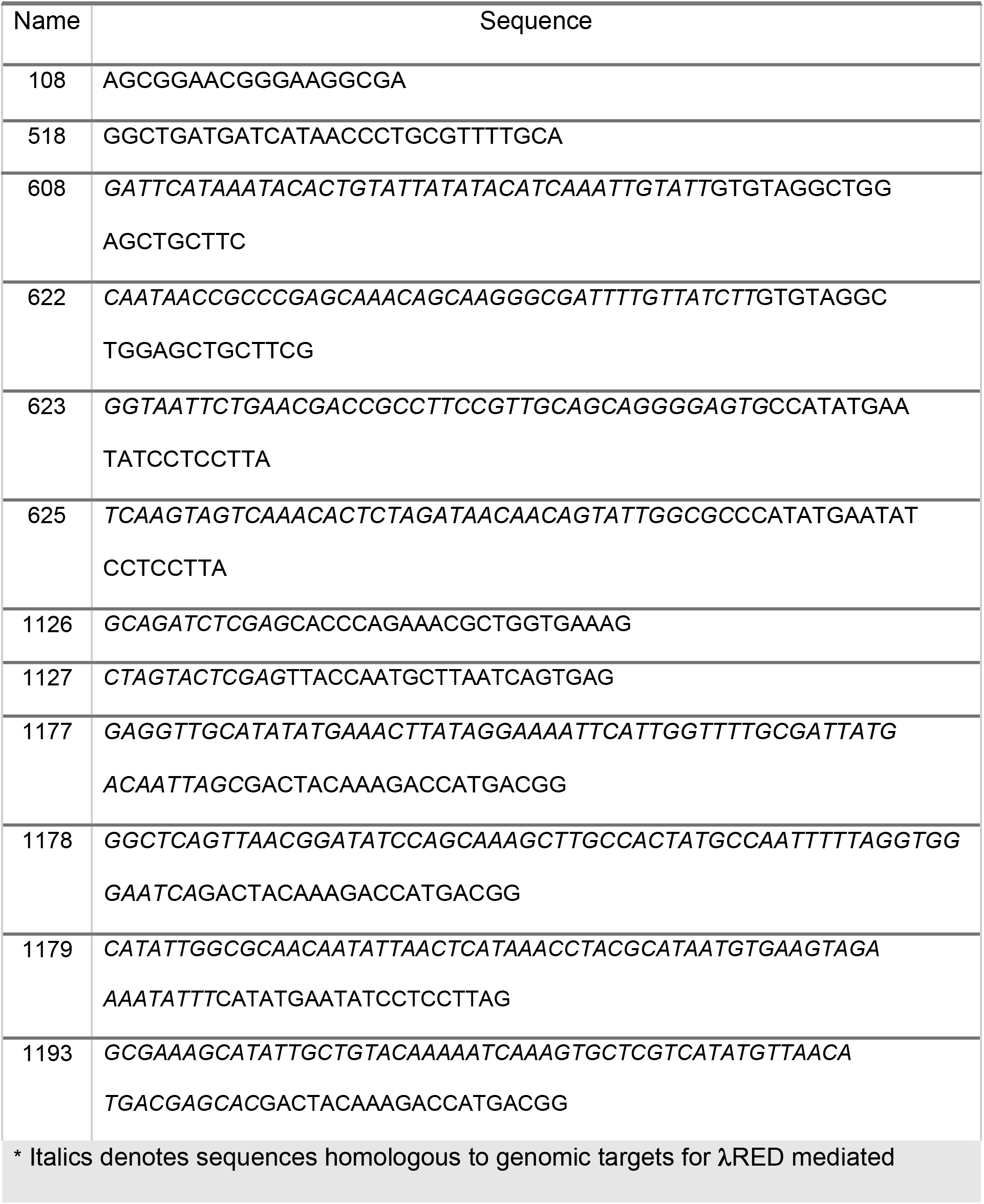
Oligonucleotide primers used in this study

### Quantitative bacterial adherence assays

HCT-8 cells, a human ileocecal epithelial cell line, was purchased from the American Type Culture Collection (ATCC CCL-244). HCT-8 cells were grown in IMDM supplemented with 10% fetal bovine serum (FBS) at 37°C in 5% CO_2_. Approximately 24 hours before infection HCT-8 cells were seeded in 24-well tissue culture plates at ca. 10^5^ cells per well. Bacteria were cultured in CFA medium to an OD_600_ of 0.7-0.8. Bacteria were collected from 500 μl of culture by centrifugation then resuspended in 1 ml IMDM, 10% FBS. The tissue culture wells were aspirated, washed once with PBS, then 200 μl IMDM,10% FBS was added to each well followed by 50 μl of the bacterial suspension for an MOI of ca. 50. After a one hour attachment period culture medium was aspirated and wells were washed four times with PBS to remove nonadherent bacteria. Adherent bacteria were collected in PBS, 0.1% (vol/vol) Triton X-100 and enumerated by serial dilution and plating. To control for differences between inocula adherent CFUs were normalized to inocula.

### Trichloroacetic acid (TCA) precipitation of culture supernatants

H10407 / pNEB193 was cultured in either CFA or IMDM to stationary phase. Bacteria were removed from the culture medium by centrifugation and successive filtration through 0.45 and 0.20 micron syringe tip filters. TCA was added to the clarified supernatants at a final concentration of 10% (vol/vol). Proteins were precipitated overnight at 4°C then pelleted for 8 min. at 21,000 x *g*. Protein pellets were washed twice with 500 μl ice-cold acetone to remove residual TCA, air-dried for 15 min. at room temperature, then resuspended in 60 μl of 1x SDS-PAGE loading buffer with 100 mM β-mercaptoethanol.

### Proteinase K digests

Bacteria were collected from 400 μl of stationary phase cultures by centrifugation at 21,000 g for 3 min. and resuspended in 75 μl of either PBS or PBS, 10 mM EDTA, 1% (vol/vol) Triton X-100. Suspensions were equilibrated at 37°C for 1 hour prior to addition of proteinase K to a final concentration of 200 μg/ml. Some samples received 11 mM PMSF prior to enzyme addition. All samples were incubated for 1 hour at 37°C with continuous gentle rocking. Enzyme activity was terminated by addition of PMSF followed by 25 μl of 4x SDS-PAGE loading buffer with β-mercaptoethanol.

### Immunoblots

Whole-cell lysates were subject to SDS-PAGE, transferred to PVDF membranes, and blocked in TBS-Blotto (25 mM TrisCl pH 7.6, 150 mM NaCl, 5% (wt/vol) powdered nonfat milk. Antibodies against CexE_α_ were produced by immunization of rabbits (Proteintech Group, Inc.) with purified CexE_α_-His6 and used at a dilution of 1:10000 in TBS-Blotto with 0.05% vol./vol. Tween20 (14). Anti-DnaK (ab69617) and anti-β-lactamase (ab12251) antibodies were purchased from AbCam and used at dilutions of 1:10000 and 1:10000 respectively. Anti-FLAG (F1804) was purchased from Sigma-Aldrich and used at a dilution of 1:10000 HRP conjugated goat anti-rabbit (sc-2030) and goat anti-mouse (115-036-062) antibodies were purchased from Santa Cruz Biotechnology and Jackson ImmunoResearch Laboratories respectively. Secondary antibodies were used at a dilutions of 1:10000. Chemiluminescence was detected with a Odyssey FC Imaging System (LI-COR Biosciences). Densitometry analysis was performed utilizing ImageJ software.

### In Vivo studies

All mice were bred and housed under barrier conditions in the Division of Veterinary Resources of the University of Miami Miller School of Medicine. Mice were regularly screened for specific common pathogens. C57Bl/6 mice were inoculated orogastrically with *C. rodentium* strains in PBS. Adults were inoculated with a 22-gauge, round-tipped feeding needle. Infant mice were used at 15 days of age and inoculated with PE-10 tubing (polyethylene tubing with an outside diameter of 0.61 mm) attached to a 30-gauge needle. Administered CFUs were determined by serial dilution and plating of inocula. Infant mice were weaned at 3 ½ weeks of age. On select days fecal pellets were collected from adult mice and homogenized in sterile ddH2O, diluted, and plated on MacConkey agar with kanamycin. Organs of adult mice were homogenized in PBS using a OMNI International Tissue Homogenizer (Kennesaw, GA) for 2 min at medium speed. Organs from infant mice were homogenized in HBSS using a Seward Biomaster 80 Stomacher (Brinkman, Westbury, NY) for 4 min. at high speed. Homogenates were diluted and plated on MacConkey agar plates with kanamycin. CFUs were normalized to the weight of fecal pellets and organs.

### Statistical analysis

One-way ANOVA with Tukey’s multiple comparisons test, Student’s *t*-test, Mann-Whitney U Test, and Mantel-Cox logrank test. (GraphPad Prism Version 7.3 was utilized for statistical analysis).

### Ethics statement

All animal experiments were approved by and performed in accordance with the University of Miami Institutional Animal Care and Use Committee guidelines.

## References

1. 2017. Diarrhoeal disease. WHO.

2. Mathers CD, Loncar D. 2006. Projections of global mortality and burden of disease from 2002 to 2030. PLoS Med 3:2011–2030.

3. Qadri F, Svennerholm A-M, Faruque ASG, Sack RB. 2005. Enterotoxigenic Escherichia coli in developing countries: epidemiology, microbiology, clinical features, treatment, and prevention. Clin Microbiol Rev 18:465–83.

4. Fleckenstein JM, Hardwidge PR, Munson GP, Rasko DA, Sommerfelt H, Steinsland H. 2010. Molecular mechanisms of enterotoxigenic Escherichia coli infection. Microbes Infect 12:89–98.

5. Evans DG, Satterwhite TK, Evans DJ, DuPont HL, DuPont HL. 1978. Differences in serological responses and excretion patterns of volunteers challenged with enterotoxigenic Escherichia coli with and without the colonization factor antigen. Infect Immun 19:883–8.

6. Kernéis S, Chauvière G, Darfeuille-Michaud A, Aubel D, Coconnier MH, Joly B, Servin AL. 1992. Expression of receptors for enterotoxigenic Escherichia coli during enterocytic differentiation of human polarized intestinal epithelial cells in culture. Infect Immun 60:2572–80.

7. Caron J, Scott JR. 1990. A rns-like regulatory gene for colonization factor antigen I (CFA/I) that controls expression of CFA/I pilin. Infect Immun 58:874–8.

8. Caron J, Coffield LM, Scott JR. 1989. A plasmid-encoded regulatory gene, rns, required for expression of the CS1 and CS2 adhesins of enterotoxigenic Escherichia coli. Proc Natl Acad Sci U S A 86:963–7.

9. Munson GP, Scott JR. 1999. Binding site recognition by Rns, a virulence regulator in the AraC family. J Bacteriol 181:2110–7.

10. Bodero MD, Harden EA, Munson GP. 2008. Transcriptional regulation of subclass 5b fimbriae. BMC Microbiol 8:180.

11. Bodero MDR, Munson GP. 2016. The Virulence Regulator Rns Activates the Expression of CS14 Pili. Genes (Basel) 7:120.

12. Crossman LC, Chaudhuri RR, Beatson SA, Wells TJ, Desvaux M, Cunningham AF, Petty NK, Mahon V, Brinkley C, Hobman JL, Savarino SJ, Turner SM, Pallen MJ, Penn CW, Parkhill J, Turner AK, Johnson TJ, Thomson NR, Smith SGJ, Henderson IR. 2010. A commensal gone bad: complete genome sequence of the prototypical enterotoxigenic Escherichia coli strain H10407. J Bacteriol 192:5822–31.

13. Porter CK, Riddle MS, Tribble DR, Louis Bougeois A, McKenzie R, Isidean SD, Sebeny P, Savarino SJ. 2011. A systematic review of experimental infections with enterotoxigenic Escherichia coli (ETEC). Vaccine 29:5869–85.

14. Pilonieta MC, Bodero MD, Munson GP. 2007. CfaD-dependent expression of a novel extracytoplasmic protein from enterotoxigenic Escherichia coli. J Bacteriol 189:5060–5067.

15. Petty NK, Bulgin R, Crepin VF, Cerdeño-Tárraga AM, Schroeder GN, Quail MA, Lennard N, Corton C, Barron A, Clark L, Toribio AL, Parkhill J, Dougan G, Frankel G, Thomson NR. 2010. The Citrobacter rodentium genome sequence reveals convergent evolution with human pathogenic Escherichia coli. J Bacteriol 192:525–38.

16. Mundy R, MacDonald TT, Dougan G, Frankel G, Wiles S. 2005. Citrobacter rodentium of mice and man. Cell Microbiol 7:1697–706.

17. Bendtsen JD, Nielsen H, von Heijne G, Brunak S. 2004. Improved prediction of signal peptides: SignalP 3.0. J Mol Biol 340:783–95.

18. Pugsley AP. 1993. The complete general secretory pathway in gram-negative bacteria. Microbiol Rev 57:50–108.

19. Kumar S, Stecher G, Tamura K. 2016. MEGA7: Molecular Evolutionary Genetics Analysis Version 7.0 for Bigger Datasets. Mol Biol Evol 33:1870–4.

20. Reuter S, Connor TR, Barquist L, Walker D, Feltwell T, Harris SR, Fookes M, Hall ME, Petty NK, Fuchs TM, Corander J, Dufour M, Ringwood T, Savin C, Bouchier C, Martin L, Miettinen M, Shubin M, Riehm JM, Laukkanen-Ninios R, Sihvonen LM, Siitonen A, Skurnik M, Falcão JP, Fukushima H, Scholz HC, Prentice MB, Wren BW, Parkhill J, Carniel E, Achtman M, McNally A, Thomson NR. 2014. Parallel independent evolution of pathogenicity within the genus Yersinia. Proc Natl Acad Sci U S A 111:6768–73.

21. Schauer DB, Falkow S. 1993. Attaching and effacing locus of a Citrobacter freundii biotype that causes transmissible murine colonic hyperplasia. Infect Immun 61:2486–92.

22. Murata T, Iida T, Shiomi Y, Tagomori K, Akeda Y, Yanagihara I, Mushiake S, Ishiguro F, Honda T. 2001. A large outbreak of foodborne infection attributed to Providencia alcalifaciens. J Infect Dis 184:1050–5.

23. Yoh M, Matsuyama J, Ohnishi M, Takagi K, Miyagi H, Mori K, Park K-S, Ono T, Honda T. 2005. Importance of Providencia species as a major cause of travellers’ diarrhoea. J Med Microbiol 54:1077–82.

24. Méric G, Hitchings MD, Pascoe B, Sheppard SK. 2016. From Escherich to the Escherichia coli genome. Lancet Infect Dis 16:634–636.

25. Khetrapal V, Mehershahi KS, Chen SL. 2017. Complete Genome Sequence of the Original Escherichia coli Isolate, Strain NCTC86. Genome Announc 5:e00243–17.

26. Sheikh J, Czeczulin JJR, Harrington S, Hicks S, Henderson IIR, Bouguénec C Le, Gounon P, Phillips AA, Nataro JJP, Le Bouguénec C, Gounon P, Phillips AA, Nataro JJP. 2002. A novel dispersin protein in enteroaggregative Escherichia coli. J Clin Invest 110:1329–1337.

27. Hart E, Yang J, Tauschek M, Kelly M, Wakefield MJ, Frankel G, Hartland EL, Robins-Browne RM. 2008. RegA, an AraC-like protein, is a global transcriptional regulator that controls virulence gene expression in Citrobacter rodentium. Infect Immun 76:5247–56.

28. Allen KP, Randolph MM, Fleckenstein JM. 2006. Importance of heat-labile enterotoxin in colonization of the adult mouse small intestine by human enterotoxigenic Escherichia coli strains. Infect Immun 74:869–75.

29. Diaz LA, Altman NH, Khan W, Serhan CN, Adkins B. 2017. Specialized Proresolving Mediators Rescue Infant Mice from Lethal Citrobacter rodentium Infection and Promote Immunity against Reinfection. Infect Immun 85.

30. Rasko DA, Rosovitz MJ, Myers GSA, Mongodin EF, Fricke WF, Gajer P, Crabtree J, Sebaihia M, Thomson NR, Chaudhuri R, Henderson IR, Sperandio V, Ravel J. 2008. The pangenome structure of Escherichia coli: comparative genomic analysis of E. coli commensal and pathogenic isolates. J Bacteriol 190:6881–93.

31. Sahl JW, Sistrunk JR, Baby NI, Begum Y, Luo Q, Sheikh A, Qadri F, Fleckenstein JM, Rasko DA. 2017. Insights into enterotoxigenic Escherichia coli diversity in Bangladesh utilizing genomic epidemiology. Sci Rep 7:3402.

32. Hazen TH, Michalski J, Luo Q, Shetty AC, Daugherty SC, Fleckenstein JM, Rasko DA. 2017. Comparative genomics and transcriptomics of Escherichia coli isolates carrying virulence factors of both enteropathogenic and enterotoxigenic E. coli.Sci Rep 7:3513.

33. Nishi J, Sheikh J, Mizuguchi K, Luisi B, Burland V, Boutin A, Rose DJ, Blattner FR, Nataro JP. 2003. The export of coat protein from enteroaggregative Escherichia coli by a specific ATP-binding cassette transporter system. J Biol Chem 278:45680–9.

34. Koronakis V, Eswaran J, Hughes C. 2004. Structure and function of TolC: the bacterial exit duct for proteins and drugs. Annu Rev Biochem 73:467–89.

35. Koronakis V, Sharff A, Koronakis E, Luisi B, Hughes C. 2000. Crystal structure of the bacterial membrane protein TolC central to multidrug efflux and protein export. Nature 405:914–9.

36. Roy K, Hamilton DJ, Munson GP, Fleckenstein JM. 2011. Outer membrane vesicles induce immune responses to virulence proteins and protect against colonization by enterotoxigenic Escherichia coli. Clin Vaccine Immunol 18:1803–1808.

37. Schwechheimer C, Kuehn MJ. 2015. Outer-membrane vesicles from Gram-negative bacteria: Biogenesis and functions. Nat Rev Microbiol.

38. Horstman AL, Kuehn MJ. 2000. Enterotoxigenic Escherichia coli secretes active heat-labile enterotoxin via outer membrane vesicles. J Biol Chem 275:12489–96.

39. Evans DG, Evans DJ, Tjoa W. 1977. Hemagglutination of human group A erythrocytes by enterotoxigenic Escherichia coli isolated from adults with diarrhea: correlation with colonization factor. Infect Immun 18:330–7.

40. Datsenko KA, Wanner BL. 2000. One-step inactivation of chromosomal genes in Escherichia coli K-12 using PCR products. Proc Natl Acad Sci U S A 97:6640–5.

41. Datta S, Costantino N, Court DL. 2006. A set of recombineering plasmids for gram-negative bacteria. Gene 379:109–15.

42. Kumar P, Luo Q, Vickers TJ, Sheikh A, Lewis WG, Fleckenstein JM. 2014. EatA, an immunogenic protective antigen of enterotoxigenic Escherichia coli, degrades intestinal mucin. Infect Immun 82:500–8.

43. Schauer DB, Zabel BA, Pedraza IF, O’Hara CM, Steigerwalt AG, Brenner DJ. 1995. Genetic and biochemical characterization of Citrobacter rodentium sp. nov.J Clin Microbiol 33:2064–8.

44. Yanisch-Perron C, Vieira J, Messing J. 1985. Improved M13 phage cloning vectors and host strains: nucleotide sequences of the M13mpl8 and pUC19 vectors. Gene 33:103–119.

45. Uzzau S, Figueroa-Bossi N, Rubino S, Bossi L. 2001. Epitope tagging of chromosomal genes in Salmonella. Proc Natl Acad Sci U S A 98:15264–9.

